# Effects of demographic stochasticity and life-history strategies on times and probabilities to fixation: an individual-based model

**DOI:** 10.1101/207100

**Authors:** Diala Abu Awad, Camille Coron

## Abstract

Previous works has suggested that the harmonic mean population size can summarize the consequences of demographic fluctuations on the genetic frequencies of populations. We test this hypothesis by studying a model in which the demography and genetic composition of the population are both determined by the behavior of the individuals within the population. We propose an effective population size that allows us to compare our model with the classical Wright-Fisher diffusion both for neutral alleles and those under selection. We find that using our approximation for the effective population size, the Wright-Fisher diffusion provides good results for the times to absorption and probabilities of fixation of a given neutral allele and in cases where selection is not too strong. However, the times and laws to fixation are not always well predicted due to large fluctuations in population size caused by small growth rates or strong competition between individuals, that cannot be captured by the constant population size approximation. The discrepancy between our model and the Wright-Fisher diffusion is accentuated in the presence of demo-genetic feed-back. Our results imply that the Wright-Fisher diffusion is not appropriate when studying probabilities and times to fixation in long-lived species with low reproductive rates.

## 1 Introduction

Adaptive and non adaptive evolution is characterized by the dynamics of allele frequencies and their eventual loss or fixation. For more than half a century, the diffusion limit of the Wright-Fisher model ([15, 37]), introduced by [22, 23], has provided one of the key tools in population genetics for predicting the dynamics of allelic frequencies. Due to simple and strong analytical results obtained for this general model ([24]), it has been extended to take into account populations with more general and complicated behaviors such as non-random mating and structured populations (see for example [2, 1, 34]). The Wright-Fisher model makes two simplifying assumptions: (1) all individuals reproduce and die at the same time (discrete non-overlapping generations), and (2) population size is fixed, which has led to the concept of “effective population size”, denoted *N*_*e*_ (and discussed below). However, population size tends to vary stochastically, notably since births and deaths can be independent events: reproduction by an individual is not necessarily immediately followed by its death (see for instance [5]), and the speeds at which reproduction and death occur representing different life-history strategies (*i.e. r/K* strategies). The ubiquity of stochastic demographic phenomena, such as extinction, rapid expansions and bottlenecks on a macroscopic scale, or independent births and deaths on a microscopic scale, requires a better understanding of their interaction with allele frequency dynamics (and notably with allele fixation).

In existing models studying allele dynamics, *N*_*e*_ is a central notion which aims at bringing any population as “close” as possible (the definition of closeness being dependent on the indicators of interest) to a classical Wright-Fisher diffusion. In particular for populations with a deterministically varying population size, this parameter is defined as the harmonic mean of the population size (as shown in [38, 25], for instance). In the presence of selection, [31] explored the impact of macroscopic demographic events (introduced by the use of a non-constant deterministic population size) on the probability of fixation of alleles. They found that the harmonic mean sufficed in reflecting the change in fixation probabilities of fluctuating populations as long as selection was not too strong. On the other hand, [21, 20] showed that the harmonic mean size is sometimes an inadequate definition of the effective population size when population size varies stochastically and the authors proposed a new definition for *N*_*e*_ (the heterozygosity effective size). The harmonic mean seems therefore insufficient in capturing the effects of stochastic events on a more microscopic level even in models where the deaths and births of individuals are not considered explicitly, the general effects of these processes being averaged to reflect the behavior of the entire population.

Recently, individual-based models examining the interaction between population size dynamics on the microscopic scale and probabilities of fixation have been developed ([5, 6, 32]). However, the feedback of genetics on demography is not considered nor modeled in these diffusions, whereas it can have a major impact on population viability, notably when selection parameters are not small, as can be observed in models of evolutionary rescue ([29, 18]), where this feedback is a central aspect. In [32], the authors explored the consequences of different life-history strategies and proposed an individual-based model with “quasineutral” selection so that the impact of population genetics on population demography can be neglected and found that they could not define an appropriate Ne for which a classical neutral Wright-Fisher diffusion would give the same mean time to absorption and fixation probability as their model. Mean times to fixation of neutral alleles, and eventually the distribution of these times, in the Wright-Fisher diffusion depend on the population’s *N*_*e*_ ([25]) and are thus expected to be affected by a population’s demographic dynamics (notably due to macroscopic events such as bottlenecks, expansions and extinctions, as can be deduced from works on coalescent theory [19]). On the contrary, the fixation probability of a neutral allele is always expected to be equal to its initial frequency. That [32]’s results for quasi-neutrality are better described by a Wright-Fisher diffusion with selection (Figure 4 in their paper) thus raises three questions: *i*) How should fitness be defined in individual-based models in order to render them, if possible, comparable to a Wright-Fisher framework? *ii*) What role do life-history strategies play in the probabilities and times to fixation? and *iii*) If genotypes under selection present different demographic behaviors (*i.e*. growth rate), how is the ensuing change in population size likely to influence the probabilities and times to fixation?

In this article we propose an individual-based model in order to study the absorption times and fixation probabilities in a demo-genetic context, which we then compare to a Wright-Fisher diffusion. In this probabilistic model both the demography and genetics of a given population are defined through the dynamics of each individual within the population. The behavior of each individual is stochastic, and dependent on demographic parameters that can be estimated ([27]). More precisely, we consider a population of diploid individuals experiencing weak selection at a single bi-allelic locus. As population size is directly determined by frequent birth and death events, it changes stochastically with time, and can also depend on the population’s genetic composition. The originality of our approach and model lies in four main features: (1) We consider linked stochastic dynamics of both the population size and its genetic composition. (2) The life-history strategy of a population is a natural behavior of the model and depends directly on the demographic parameters (as in *e.g*. [32]), being in no way forced. (3) Extinction occurs in finite time, which notably impacts fixation times. (4) We consider a sexually reproducing diploid population, with general dominance relationships between alleles, and possibility of self-fertilization (previous models considered haploid individuals, [5, 6, 32]). The obtained model can also be seen as a generalization of the Wright-Fisher diffusion, since this diffusion can be obtained when letting some parameters of the model (namely the growth and competition rates) go to +∞. We compare the laws of the time to absorption (fixation or loss of a given allele) and the probability of fixation for our model to the classical Wright-Fisher diffusion (presented in [2] for populations with self-fertilization). These results are obtained by simulating trajectories of diffusion processes, and we find notably that

i. There are parameter sets for which the laws of the time to absorption for our demo-genetic model and for the classical population genetics model (Wright-Fisher diffusion calibrated with an appropriate effective size) are very close, notably when there is a high population growth rate and high death rate (due to competition for resources). Population genetics models therefore provide very good predictions for species with *r*-strategies (high reproductive output and short life-span).
ii. Laws of time to absorption can be very different when taking into account population size variability, notably in rapid expansion and diminution contexts (Section 3.3), as well as when population size fluctuations are highly stochastic, which is the case for populations with low reproductive rates and low death rates (*K-strategies*). In particular, we find that due to the fluctuations in population size, the frequencies of small and large absorption times of rare alleles are underestimated in the classical Wright-Fisher diffusion model (*i.e.* there is a greater variance in times to absorption than predicted by a fixed population size).
iii. The demographic consequences of taking the feedback of genetics on demography can impact the probabilities and times to fixation in a way that can not be fully captured by the proposed effective population size if population growth rates are low.

## 2 Model

We consider a population of diploid individuals, characterized by their genotype at a single bi-allelic locus with alleles *A* and *a*. The population is modeled by a 3-dimensional stochastic birth-and-death process (detailed below) giving the respective numbers of individuals with genotype *AA*, *Aa* and *aa*. Contrary to previous models where population size is a parameter, here it is a random variable. The dynamics of population size are stochastic, and population extinction occurs with probability 1. Below we detail the rescaled diffusion approximation, highlighting the main differences between our model and the diffusion approximation proposed by [24].

### 2.1 Rescaled birth-and-death process

The Wright-Fisher diffusion is obtained by considering the dynamics of the proportion of a given allele when re-scaling a discrete time population model with constant effective population size *N*_*e*_ and non-overlapping generations. In this article, population dynamics are determined by individual-based demographic parameters, therefore inducing variable population size. We introduce a scaling parameter *K* ∈ {1, 2,…} that will go to infinity (as in [16, 7, 9, 10]), modeling an infinite size approximation. The population is made up of three types of individuals (*AA*, *Aa* and *aa*, represented by 1, 2 and 3 respectively), the number of individuals of each type being of order *K*. At each time *t* the population is represented by a vector

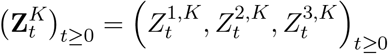

which gives the respective number of individuals of each type, divided by *K*. If the population is at a state **z** = (*z*_1_,*z*_2_,*z*_3_), the birth rates 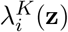 for all *i* ∈ {1, 2,3} model sexual Mendelian reproduction either by self-fertilization (with probability *α*) or by random mating (with probability 1 − *α*).

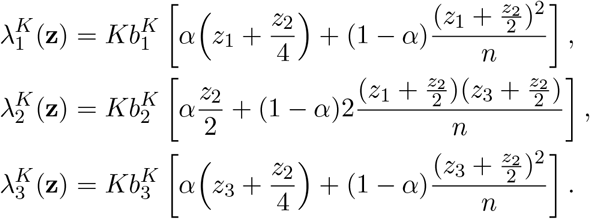

with *n* = *z*_1_+*z*_2_+*z*_3_ ≠ 0. These birth rates are naturally set to 0 when *n* = 0. Note that the parameters 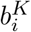 that model the viability (or recruitment) of new-born individuals can depend on *i*, which allows for some selection at birth (as will be shown below). Individual mortality can be natural or due to competition with other individuals (therefore allowing for density-dependence and limiting population size). Here we assume that death rates do not depend on genotypes, in order to focus on a small number of parameters (but see [11] for a more general model). If the population is at a state **z** = (*z*_1_,*z*_2_,*z*_3_, the rate 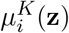 at which an individual with genotype i dies in the population is then given by:

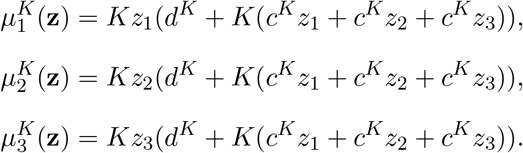

The demographic parameter *d*^*K*^ (resp. *c*^*K*^ > 0) is the intrinsic death rate (resp. the competition rate) of individuals. Population size is therefore regulated by competition, *i.e*. by density-dependence.

The demographic parameters *b*^*K*^, *d*^*K*^ and *c*^*K*^ are scaled both by *K* and a parameter *γ*, the latter scaling the speed with which births and deaths occur, giving:

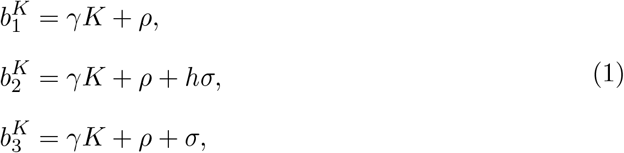

and

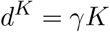 and 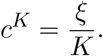

The parameters *σ* and *h* are respectively the selection and dominance coefficients of allele *a*, and *ρ* is the population growth rate in the absence of selection. Note that in this model, we do not directly consider population size (number of individuals), but population mass, defined as 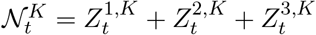. This scaling therefore models a population with numerous and small individuals, each represented by a mass of 1/*K*, that reproduce frequently in such a way that both the total population mass and allele proportions will not be constant (and will evolve stochastically) even when the scaling parameter *K* goes to infinity. This is the same scaling used to obtain the Wright-Fisher diffusion process from the Wright-Fisher model, however our initial model (birth-and-death process) allows for stochastic dynamics of population mass. When *K* is large, the selection parameter *σ* has an inherent weak impact on the birth parameters 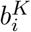, but due to first-order compensation between birth and death events (both of order *K*), its impact on the growth rate is macroscopic. Therefore, it will still have an effect on the limiting population dynamics (notably by either increasing or decreasing the expected population mass, see next section).

### 2.2 Extended Hardy-Weinberg structure and limiting diffusion process

Let us set for all *K* ≥ 1 and all *t* ≥ 0,

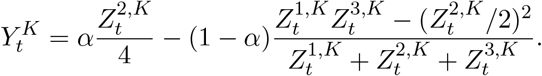

Note that in a pure random mating context (*α* = 0), and if the quantity 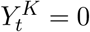, then the proportion of each genotype in the population is equal to the proportion of pairs of alleles forming this genotype, which means that the population satisfies the Hardy-Weinberg structure. More generally, 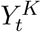 quantifies the deviation of the population at time *t* from a generalized Hardy-Weinberg structure. Indeed, straightforward calculations show that, as in population genetics theory ([17], pp. 91-93), if 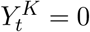 then

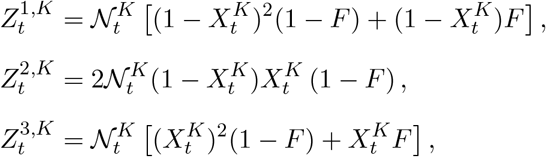

where 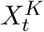 is the proportion of allele *a* in the population, and the coefficient of inbreeding 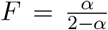. We can prove following [10] that 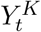 converges to 0 when *K* goes to infinity, for all *t*. The limiting population dynamics can then be represented at time *t* by the couple 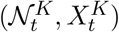 giving the population mass and the proportion of allele *a*. The population process 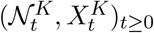 thus converges toward a bi-dimensional diffusion process 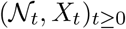 whose equation can be written as:

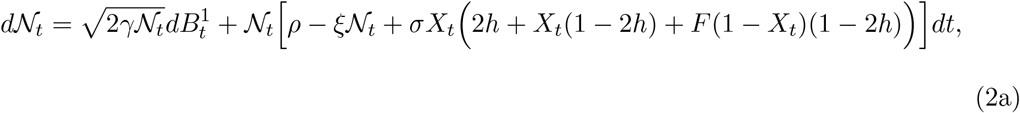

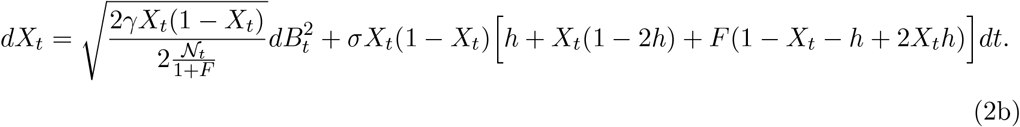

where 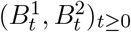 is a bi-dimensional standard Brownian motion (stochastic component of the equation). This diffusion model can be generalized without difficulty to any finite number of alleles, as presented in [12]. Note that, without loss of generality, we can assume that the time scaling parameter *γ* is equal to 1/2, thus simplifying the above equations. In this case, if the stochastic quantity 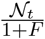 is artificially replaced by a fixed parameter *N*_*e*_, then the model given in (2b) is the Wright-Fisher diffusion with selection and selffertilization presented in [2], where the parameter *σ* in our model is equal to the coefficient of selection *s* of [2] and *N*_*e*_ is the effective population mass.

More interestingly, this classical Wright-Fisher diffusion with selection and self-fertilization can also be directly retrieved from our model, by setting *ρ/ξ* = *N*_*e*_ and letting *ρ* got to +∞.In order to determine whether a constant effective population mass can summarize the effects of a stochastic population mass as proposed in earlier models [24, 31], in Section 33.2 we define a fixed effective population mass *N*_*e*_ in such a way that the model in [2] is adequately calibrated.

### 2.3 Simulating the diffusion process

In [2], the authors provide explicit formulas for the probabilities of fixation as well as approximations for the times to loss or fixation of an allele. Due to the bi-dimensionality of our model which largely increases the difficulty of mathematical calculations, fixation probabilities as well as laws of times to fixation, loss and absorption (either loss or fixation) of allele *a* are determined using simulations of equations (2a) and (2b). These simulations are run using a script written in C++ (and available on Dryad). The stochastic elements of the equations, 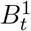 and 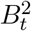 are obtained by successive samplings from a normal distribution with mean 0 and variance *dt*. *dt* is the size of the time step and is a parameter fixed at the beginning of the simulation, which we have set to 10^−4^ for a carrying capacity 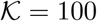 and and 10^−5^ for 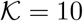 and 1. Each simulation is run until the allele *a* is either lost or fixed and 100 thousand replicas are run for each parameter set (unless otherwise mentioned) from which the probability of fixation, as well as the means and laws of times to fixation, loss and absorption are obtained.

In order to test whether deviations in times to loss or fixation from the approximations provided in [2] are due to demographic stochasticity or due to the approximations made, we run simulations of the Wright-Fisher Diffusion (using a fixed population mass *N*_*e*_ defined in Section 33.2). We also run simulations to assess the effects of the feed-back between selection and demography by artificially setting *σ* = 0 in Equation (2a) only. In order to evaluate the effect of the change in population size due to the fixation of an allele under selection with an effect σ, we also consider the case where the carrying capacity is equal to (*ρ* + *σ*)/*ξ* (see Section 3.4).

## 3 Analytical and numerical results

### 3.1 Demography

The change in population mass given in Equation (2a) is made up of a stochastic term (dependent on 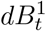) and a deterministic one (dependent on *dt*). In this diffusion model with selection and self-fertilization, the probability of extinction is equal to 1. The law of the time to extinction depends on the ecological and genetic parameters. In the neutral case where *σ* = 0, Equation (2) can be simplified the following way:

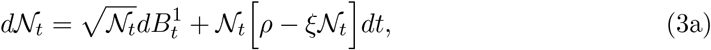

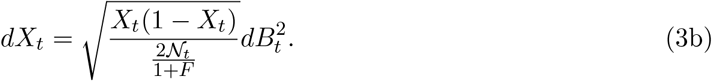

Here population mass is independent of its genetic composition and the deterministic term of Equation (3a) cancels out when 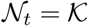 where

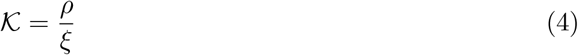

is defined as the population’s carrying capacity. Note that 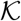 does not represent the number of individuals that can be sustained in the population (since 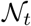 is scaled by *K* which goes to infinity) but is an indicator of the amplitude of demographic stochasticity, as will be shown below. When population mass is smaller (resp. larger) than 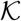, it will tend to increase (resp. decrease). For a fixed value of 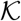, if *ρ* is large, then the population mass will remain close to 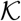, whereas for small values of ρ the mass will tend to deviate further from 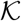 (see Figure 1). The smaller *ρ* the slower the population mass will come back to its pseudo equilibrium 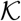; therefore a small value of *ρ* can have an important impact on extinction, as can be seen in Figure 1 (black lines). The role of 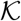 on population mass dynamics is not as straightforward since 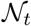 is implicated in both the stochastic and deterministic terms (therefore both terms are increased when 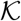 increases). In Figure 1 we also see that the effect of *ρ* on demographic stochasticity is weaker when 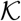 is smaller.

### 3.2 Effective population mass

In the neutral case (Equation (3)), variations in population mass are modeled by a logistic diffusion process (and thus are independent from the genetic composition of the population) and changes in allele frequency by a Wright-Fisher diffusion with population mass 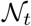 at any time *t*. Hence, it is natural to compare this model to the neutral Wright-Fisher diffusion model of population genetics, for which the proportion 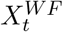 of a neutral allele at all time satisfies

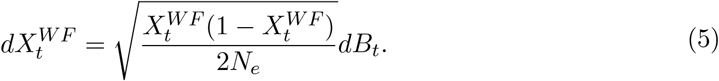

**Figure 1:**
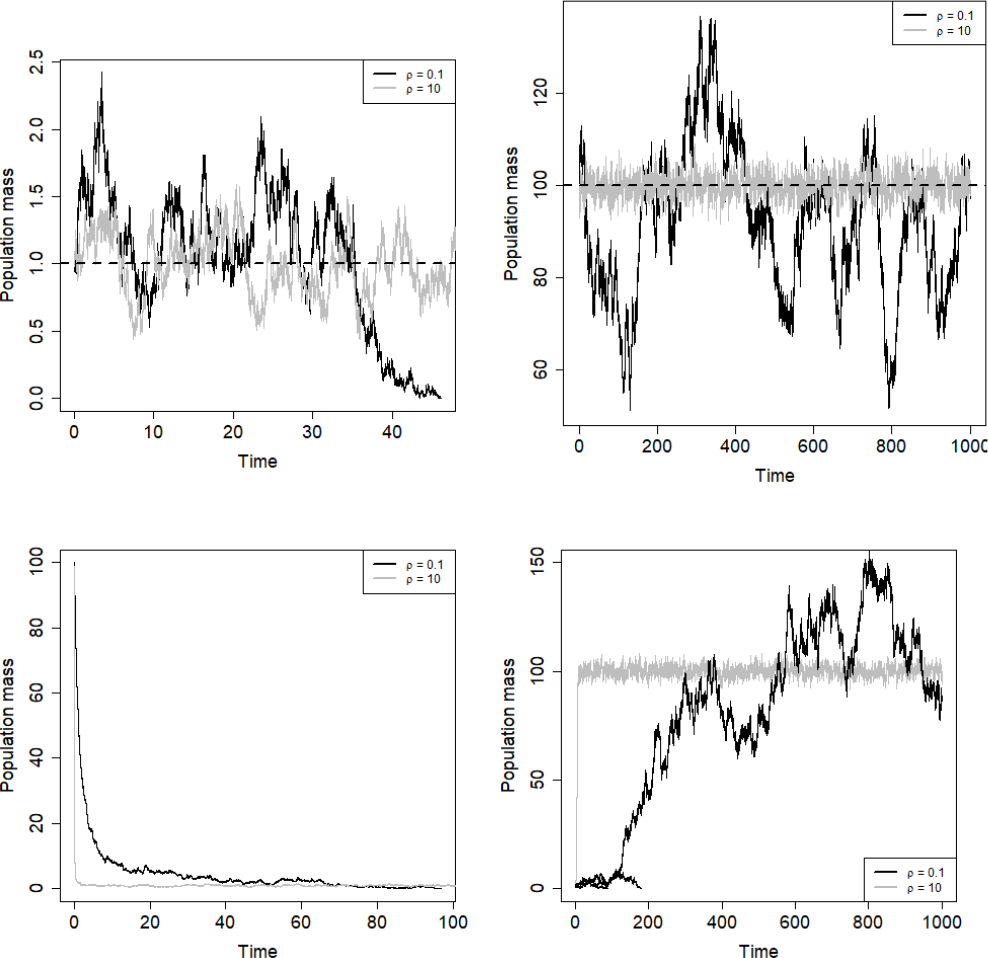
Top: Trajectories of the population mass (*N*_*t*_,*t* ≥ 0), for 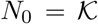 and 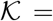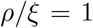 (left), 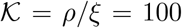 (right), for *ρ* = 0.1 (black) and *ρ* = 10 (grey). Bottom: Trajectories of the population mass (*N*_*t*_,*t* ≥ 0), for 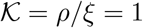 and *N*_0_ = 100 (left), and 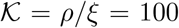 and *N*_0_ = 1 (right), for *ρ* = 0.1 (black) and *ρ* =10 (grey). For *N*_0_ = 1, *ρ* = 0.1 and 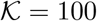 (bottom-right figure), we plot 3 trajectories.

Here *N*_*e*_ represents the effective population mass of a self-fertilizing population (as described in [2]) and is a parameter of the Wright-Fisher diffusion model. The parameters *ρ*, *ξ* and the inbreeding coefficient *F* being fixed in our model, we define a fixed effective population mass *N*_*e*_ that allows us to compare our model with variable population mass to a Wright-Fisher diffusion.

In order to calibrate *N*_*e*_ appropriately, it is not enough for the probability of fixation to be the same in both models, as in the neutral case the fixation probability of an allele *a* is simply equal to its initial proportion. Therefore, we choose to calibrate *N_e_* such that the mean absorption time (mean time to fixation of one of the two alleles) is the same in both models. From Appendix A, *N*_*e*_ is defined as:

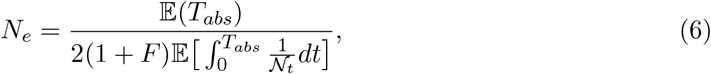

where 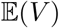 represents the expectation of a stochastic variable *V* and *T*_*abs*_ the random absorption time of the population modeled by Equation (3). Note that 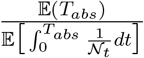 is not the expectation of the empirical harmonic mean of the mass till absorption, which is 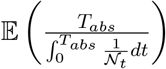, but the ratio of two expectations (the difference between the two is shown in Figure A.1, and is very important for highly fluctuating population mass). Note also that with this definition, the effective population mass *N*_*e*_ depends on the initial frequency *X*_0_ of allele a; this dependence is illustrated in Figure A.1. We obtain numerical estimations of the quantity *N*_*e*_ from the simulation runs of Equation (2) with varying population mass, calculating 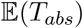 and 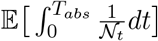 using all repetitions run for each parameter set. In Figure 2 (left) we plot the mean times to absorption as a function of the initial proportion of allele *a*, and for different values of *ρ*. This mean time to absorption is given for our model with varying population mass, for the Wright-Fisher diffusion (5) using the effective population mass *N*_*e*_ given in Equation (6), as well the theoretical result provided in Equations (12) and (13) from [2]. Figure 2 therefore shows that the models are indeed correctly calibrated for different values of parameters *ρ*, *ξ* and *X*_0_ (for different population densities and the effect of the inbreeding coefficient *F* see Figure A.2).

### 3.3 Neutral case: absorption and fixation times laws

Despite equal mean absorption times, the distributions of the times to absorption differ between our model with stochastically varying population mass and the simulation runs of the Wright-Fisher diffusion, notably when the parameters *ρ* and 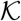 are small. This is illustrated by Figure 3 in which we compare the variance of the time to absorption for our demogenetic model and for the Wright-Fisher diffusion (see also Supplementary Figure 1, in which the laws of these times to absorption are given), and this can be understood by decomposing the absorption time into the time to loss or time to fixation of an allele at initial frequency *X*_0_. Indeed we find that mean fixation times of minority alleles are lower for the model with stochastically varying population mass (Figure 2 (right) and Supplementary Figure 2). This discrepancy between the results with varying and fixed sizes can be explained by the incidence of bottlenecks and extinction events, which is further accentuated by a small value of *ρ*. This is because a low growth rate results in a weaker impact of the deterministic forces regulating population mass (Equation (3)), further increasing demographic stochasticity. Indeed, large demographic fluctuations eventually lead to reduced population mass harmonic means, for which absorption is more rapid and fixation of minority alleles is favored (Figure 4).

**Figure 2:**
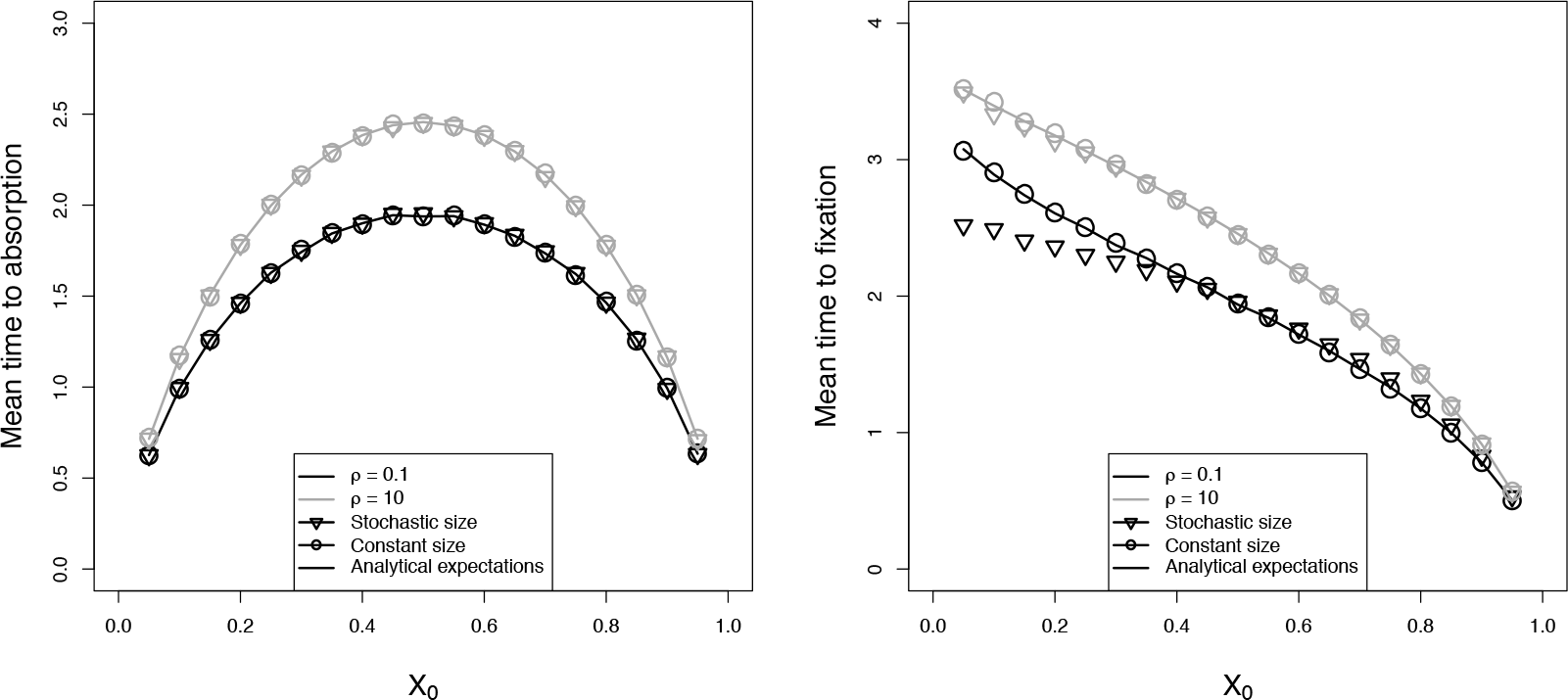
Mean times to absorption (left) and fixation (right) of a neutral allele (*σ* = 0) as a function of the initial frequency *X*_0_ of allele *a*, for three cases: 1) Simulations of the stochastic diffusion process (2) (squares), 2) Simulations of the Wright-Fisher diffusion using *N*_*e*_ defined in Equation (6) (circles) and 3) Theoretical approximations provided by [2] using *N*_*e*_ (triangles). Here we considered pure random mating (*α* = 0), the carrying capacity 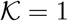 and the growth rate ρ equals 0.1 (black) or 10 (grey).

As seen in Section 3.1, we can also consider that population mass changes drastically with time, allowing us to modeling founder effects, or drastic changes in the environment for instance. As previously, we compare the laws of the absorption time in populations with rapidly decreasing or increasing mass. Population mass trajectories are given in Section 3.1 (Figure 1 (bottom)), and we start with a proportion *X* = 0.1 of a neutral allele *a*. We obtain that the laws of the absorption and fixation times are very different when comparing our to the Wright-Fisher diffusion model, despite the same mean absorption times (Figure 5). In particular, when population mass is kept constant, the frequency of small (and relatively large) absorption times is underestimated when the population mass increases, whereas the opposite is true when the population mass decreases.

**Figure 3:**
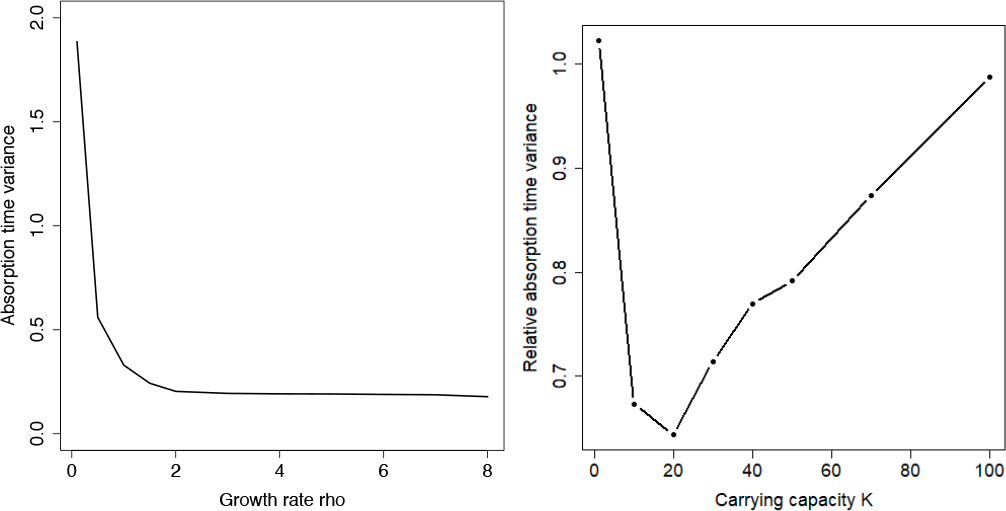
Variance of the absorption time in our demogenetic model and for the Wright-Fisher diffusion model, as a function of growth parameter *ρ* (left), and carrying capacity 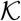(right). On the left 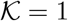 while on the right *ρ* = 0.1.

**Figure 4:**
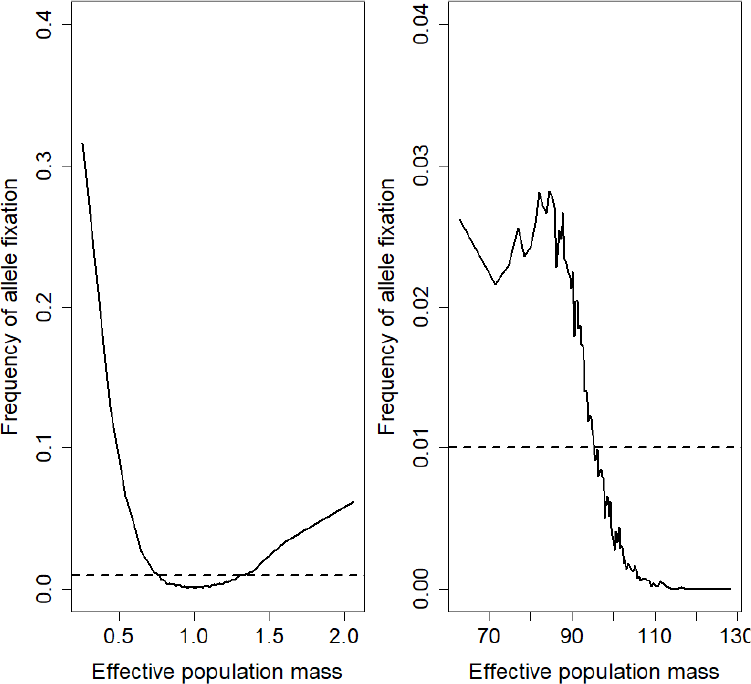
Fixation probability of a rare neutral allele, as a function of effective population mass. We set *X*_0_ = 0.01 and *ρ* = 0.1. On the left, 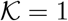 (*ξ* = 0.1), while on the right 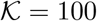(*ξ* = 0.001).

### 3.4 Selection, demography and genetic feedback

In this section we introduce selection through the parameter *σ* in Equation (2). As mentioned in Section 2.2, when comparing (2b) which describes the dynamics of allelic frequencies, to the Wright-Fisher diffusion, we find that *σ* has the same effect on allelic frequencies as the conventionally used coefficient of selection *s*. The presence of *σ* in Equation (2a) implies that population mass and the proportion of allele *a* are linked through the dynamics of individuals that are present in the population. It is important to note that, from Equation (1), selection is in fact weak and has a negligible impact on individual birth rates (whatever value of the selection parameter *σ* ∈ ℝ). However, the proportion of a given non-neutral allele can have an important impact on the population mass dynamics. The consequences of this interaction can be quantified by the ratio *σ*/*ξ*, which is the change in the carrying capacity 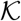 when the allele under selection *a* is fixed. Therefore, for a same 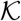 before fixation but different values of *ρ*, similar values of *σ* can lead to very different population mass dynamics (see Figure 6 with selection for a beneficial allele).

**Figure 5:**
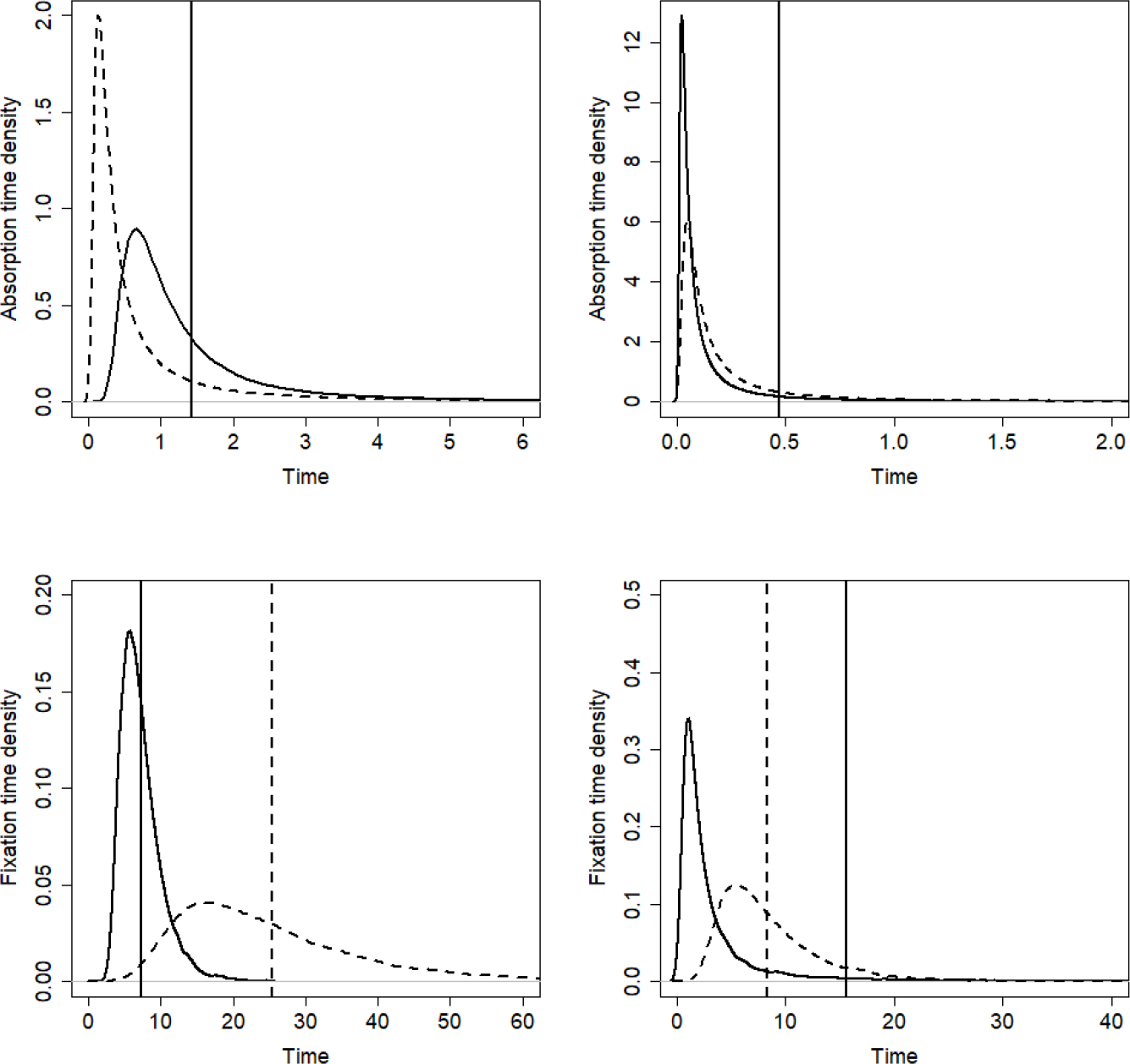
Absorption (top) and fixation (bottom) time density of a neutral allele with initial frequency *X*_0_ = 0.01 for our model and for the classical Wright-Fisher model (dotted line). On the left (decreasing population mass), we fix *N*_0_ = 100 and 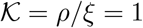, while on the right (increasing population mass), we fix *N*_0_ = 1 and 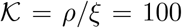, with *ρ* = 0.1.

**Figure 6:**
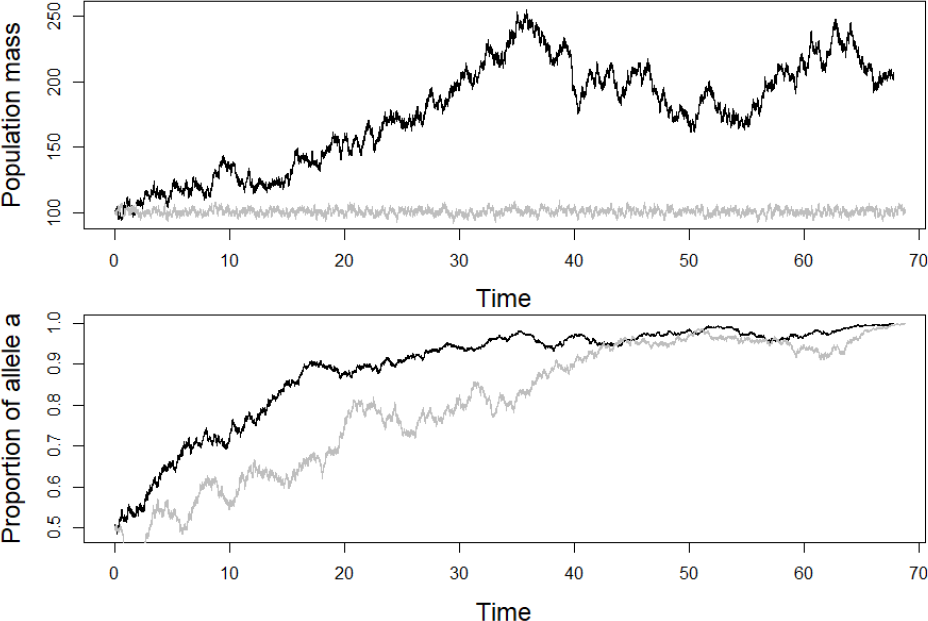
Population mass and proportion of allele *a* dynamics, for *σ* = 0.1, 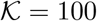, with *ρ* = 0.1 (black, 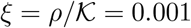) and *ρ* = 10 (gray, 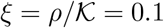).

**Figure 7:**
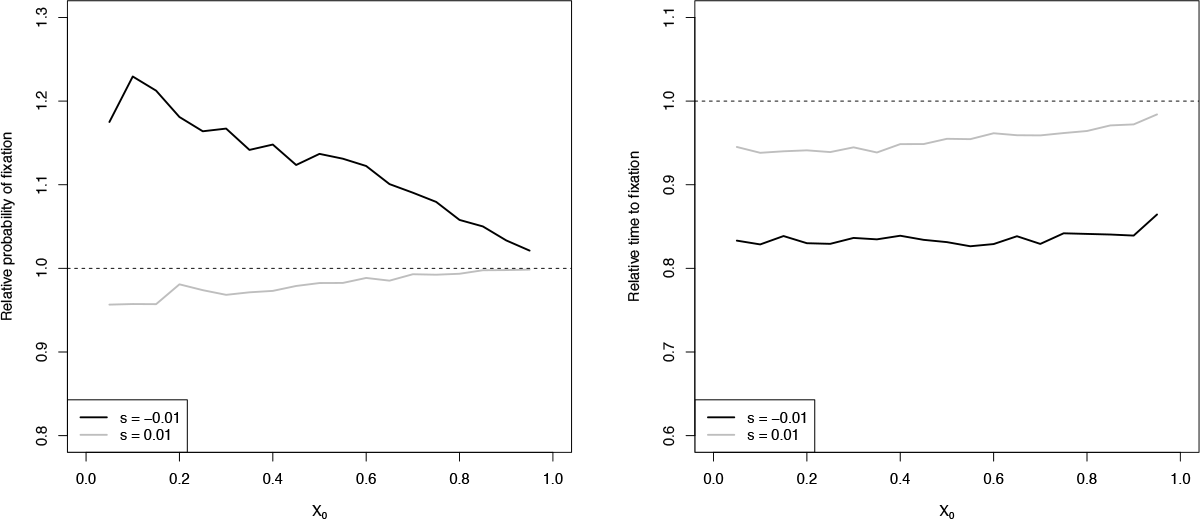
Relative probability of fixation (left) and relative time to fixation (right) for low growth rate (*ρ* = 0.1) compared to high growth rate (*ρ* = 10) as a function of the initial frequency *X*_0_ of allele *a* with 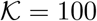, *h* = 0.25 and *α* = 0 for *s* = 0.01 and −0.01.

Due to the differences in population dynamics, probabilities and times to fixation can be affected by the growth rate, even for small values of *s* (Figure 7). Lower *ρ* results in higher probabilities of fixation of deleterious alleles, and lower relative probabilities of fixing beneficial alleles. Furthermore, times to fixation are generally lower for populations with low growth rates, independently of the coefficient of selection.

In order to understand and quantify the consequences of feedback of genetics on demography, it is natural to artificially remove all terms dependent on *σ* in Equation (2a), hence removing any impact of changes in proportion on the dynamics of population mass. More precisely, let us for simplicity assume that *F* = 0, *h* = 1/2, and let us consider the following diffusion process 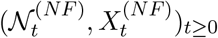 (”NF” standing for “No Feed-back”):

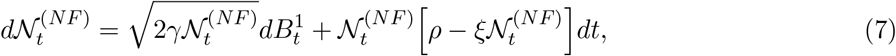

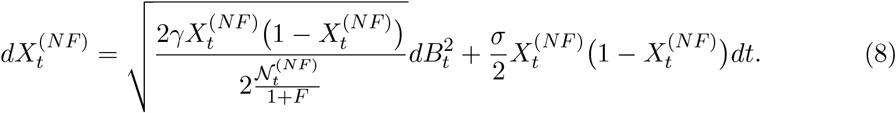

For this model without feedback, we obtain that it is possible to calibrate a Wright-Fisher diffusion with selection, using Equation (6) with 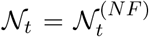, so that the mean time to absorption and the probability of fixation are the same in both models (Figure 8). In the presence of feed-back (Equation (2)), though we generally find that for large 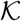, large *ρ* and/or weak selection, the proposed *N*_*e*_ (Equation (5)) provides a good approximation for the demographic effects on the times and probabilities of fixation, this is not the case for small values of *ρ* and/or 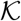. Indeed, when *ρ* is small there can be some discrepancies between the probability of fixation predicted by our model with feed-back and a population with constant size *N*_*e*_ when selection is intermediate. This can be seen in Figure 8 for *s* = 0.1 where our model with feed-back predicts a probability of up to 10% lower than the population with constant size *N*_*e*_ for low initial frequencies of the allele *a*. This difference is even greater for deleterious alleles with *s* = −0.1 (but for intermediate initial frequencies), simultaneously due to the stochastic nature of population mass and to feedback which further contributes to decreasing the population mass in this case (Figure 8). Times to fixation however are well predicted, with generally the model with feedback being either closer to the model without feed-back and 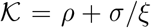 and 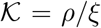 depending on the initial frequency of the allele *X*_0_. We can see from the densities of times to absorption, fixation and loss (Supplementary Figure 3), that the laws of the times to fixation are very well captured using the constant mass model, though the times to loss are slightly underestimated. Times to fixation of a mildly deleterious allele are however slightly underestimated by the simulations run with constant mass and are closer to the times to fixation of the simulations run without feed-back and 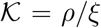.

**Figure 8:**
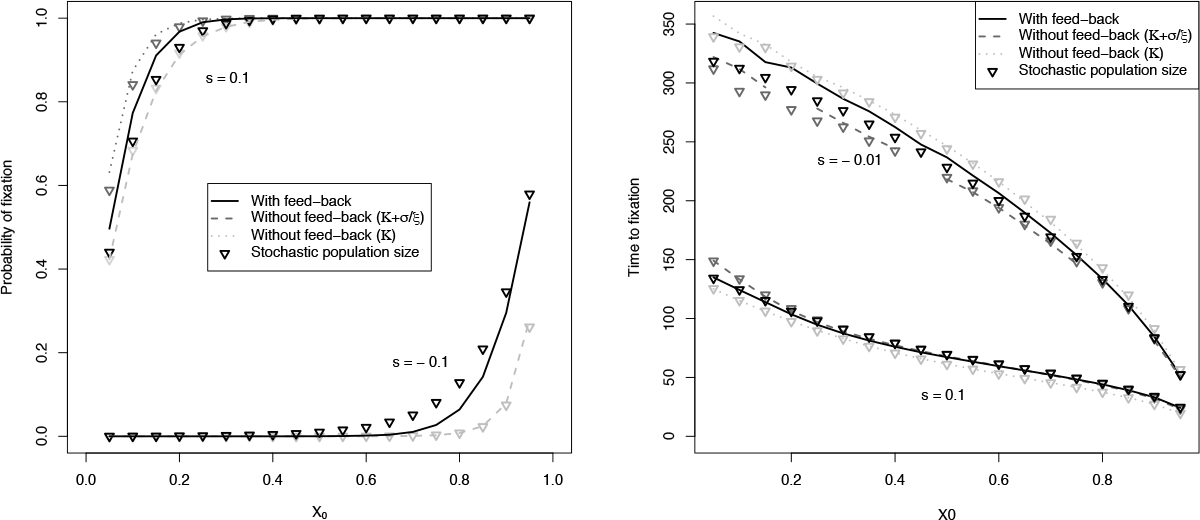
Left: Probability of fixation as a function of the initial frequency *X*_0_ of allele *a* with 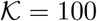, *ρ* = 0.1 for *s* = 0.1 and −0.1 from simulations with demo-genetic feedback (black full lines) and without demo-genetic feed-back (for 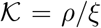 (dotted lines) and 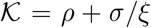 (dashed lines, not shown for *s* = −0.1 as 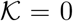 in this case), and the corresponding results of simulations run with constant mass using the corresponding *N*_*e*_. Other parameter values : *h* = 0.25 and *α* = 0. Right: Time to fixation of an allele under selection as a function of its initial frequency, same parameters as the figure on the left but for *s* = 0.1 and −0.01

Concerning the effect of the self-fertilization rate (which are summarized in Supplementary Figure 4) we find that as expected from the Wright Fisher diffusion, probabilities of fixation of beneficial (respectively deleterious) alleles increase (respectively decrease) with the rate of self-fertilization *α*. We also find as previously predicted that the times to fixation decrease with increasing *α*. In all other aspects we find the same patterns as for the case without self-fertilization (*α* = 0).

## 4 Discussion

An interesting feature of our model is that it is individual-based, in the sense that the model is characterized by simple demographic parameters that define the behavior of individuals within the population. Using these demographic parameters we are able to calculate an effective population mass *N*_*e*_ that allows us to predict the probabilities of fixation, as well as the times to absorption, using a Wright-Fisher diffusion and specify for which parameter sets this *N*_*e*_ is appropriate. We generally find that for populations with long-term fluctuations, induced by their intrinsic demographic parameters, the proposed *N*_*e*_ does not fully capture the laws of times to fixation, with rare neutral alleles being more frequently fixed in shorter times. We also show that, contrary to expectations, despite a probability of fixation of a neutral allele being equal to its initial frequency, when examining each repetition for a given parameter set separately, there is a higher frequency of fixation of rare neutral alleles for populations that maintain low harmonic mean masses. This result further highlights the importance of integrating demographic parameters in population genetics models.

### 4.1 Interpreting demographic parameters

In our model the term *ρ* defines the speed at which individuals reproduce (hence population growth) and *ξ* represents the competition for resources that in turn regulates population mass (due to increased mortality). Thus, for a given expected population mass 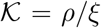 a low *ρ* describes long-lived individuals with low death rates, whereas a high *ρ* describes short-lived individuals with high death rates (rapid turnover). When comparing the demographic fluctuations of two populations with different values of *ρ*, the short-term and long-term fluctuations observed for low *ρ* and very rapid short-term fluctuations for high *ρ* (Figure 1) agree with the patterns observed for long- and short-lived species respectively (Figure 1.1 in [26]). For a same 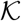 we estimate a lower *N*_*e*_ for long-lived species simultaneously due to larger population fluctuations and to the differences in population turnover speeds (since for low 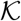 both high *ρ* and low *ρ* populations have similar fluctuations and yet we observe lower expected *N*_*e*_), which implies that on the long run a population with low *ρ* would be expected to maintain lower diversity. This prediction is supported by the lower than expected times to fixation of both neutral alleles and those under selection, as well as higher fixation probabilities of deleterious alleles for low *ρ* (see Figure 7), which agrees with the observation of less efficient purifying selection in long-lived species with low reproductive rates compared to that of short-lived ones with high reproductive rates[33, 8]. Indeed, our results indicate that in a stable environment, the stochastic demographic fluctuations and the differences in the turnover speeds of species with differing *r/K* life strategies may suffice in explaining these observations. This could explain why [33, 8] found that past historical demographic disturbances were less explicative than life-history strategies concerning contemporary genetic diversity.

### 4.2 Defining selection and fitness

One of the difficulties brought by individual-based models is how to define fitness so that it remains compatible with existing population genetics models. Indeed, several definitions of fitness do exist in literature (reviewed in [13, 30]), fitness generally being defined as a measure of the contribution of a given entity (allele, group of alleles, individual, …) to the next generation, but the notion of generation in an individual-based model is not obvious. A first way to define fitness is to focus on the Wrightian fitness (see [39]), which is defined as the mean number of progeny per individual. In the logistic birth-and-death model introduced in Section 2, the expected number of offspring for an individual with reproduction rate *b*, natural death rate *d* and competition death rate *c* in a population with (let us say fixed for simplicity) size *N* is equal to *b*/(*d* + *cN*). Obviously, when a population is at its demographic equilibrium *N* = (*b* − *d*)/*c* where births and deaths compensate, the fitness of each individual is equal to 1. In this framework the effect of a non-neutral allele or genotype (*i.e*. its coefficient of selection) can be defined as *b*′/(*d*′ + *c*′*N*) − *b*/(*d* + *cN*) = 0 if *b*′/*b* = *c*′/*c* = *d*′/*d* (where *b′*, *c*′ and *d′* respectively represent the new genotype’s birth competition and death rate). However, as shown by the results obtained for “quasi-neutral” selection in [32], where genotypes with the same Wrightian fitness but different values of *b* were considered, this definition is not sufficient in a continuous time frame. Hence a second way to consider fitness is to focus on the Malthusian fitness, which is defined as the growth rate of the population size. With this definition, fitness for our logistic birth-and-death model can be defined by the quantity [*b*/(*d* + *cN*)] × (*b* + *d* + *cN*) = *W* × *V* where *W* is the Wrightian fitness and *V* measures the speed of reproduction and death of individuals. This second definition of fitness is a more appropriate definition of fitness when studying differences in life-history strategies, as done in [32]. For both of these definitions, fitness is a quantity that is not inherent to the individual but depends on one side on demographic parameters and on the other side on both the population size and, in a non neutral framework, its genetic composition. This releases the exponential growth hypothesis naturally emerging from a concept of constant individual absolute fitness ([29]).

In this present work, we have chosen to take into account only the Wrightian fitness so as to first explore the consequences of demographic stochasticity in a model with the same genetic properties as the Wright-Fisher diffusion. Our main conclusion is that, depending on the life-history strategy of a population, the Wright-Fisher diffusion is not always able to capture the trajectories of allelic frequencies. Future work on defining an expression for the coefficient of selection in which the speed of reproduction and death *V* is also included may provide a better bridge between individual-centered models and the more mathematically manipulable Wright-Fisher diffusion.

### 4.3 Implications for empirical works

Various methods have been developed to estimate the effective size of populations (see [35] and references therein) with the aim of understanding their past and, in some cases, predicting their future evolution. However, contemporary genetic data can be greatly affected by historical events and so *N*_*e*_ is a parameter that is very population dependent ([36]). Furthermore, from an experimental point of view, the intricacy of population dynamics and population genetics requires the definition of theoretical models whose parameters can be estimated using laboratory experiments for a better understanding of their respective behaviors (reviewed in [28]). Here we provide another definition for *N*_*e*_ that is a result of both the demographic parameters of a population and, in the case of selection, its genetic properties. We find, that contrary to previous works the effects of demographic fluctuations can not always be summarized using the mean harmonic population size as proposed in [14, 24, 31]. Using the harmonic mean size is valid only when population fluctuations are sufficiently fast compared to the coalescent times [35], hence for populations with a large growth rate *ρ* and high death rates due to competition (parameter *ξ*), which represent short-lived species with high reproductive rates. This remains true even for strong fluctuations in population size when the carrying capacity 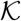 in low. However, for long lived species times to fixation cannot be summarized by *N*_*e*_, this being greatly due to near extinction events, often ignored in deterministic models (see Chapter 1 in [26]), that can contribute to lower times to fixation. Thus depending on life-history and population size, the Wright-Fisher diffusion is more or less appropriate in predicting population evolution. Though maintained genetic polymorphism is often used as a proxy for adaptive potential, one can also argue that the speed at which an advantageous allele goes to fixation is also important, especially in the face of environmental change ([18] REF). According to our model, long lived species will have a tendency to have lower probabilities of fixation of advantageous alleles, but this may be compensated by the speed at which this fixation occurs compared to that observed in short-lived species.

Previous works on integrating stochasticity into demographic models have done so by introducing a demographic variance, meant to reflect the differences between individuals in their survival and reproduction, into deterministic models (see for example [26]). However, as [26] point out, empirical measures of demographic variance may be difficult to obtain, all the more so in the ubiquitous presence of environmental stochasticity. One of the properties of our proposed models is that inter-individual variance occurs naturally, depending on the death and birth rates, and very few parameters are required in order for this variance to be ensured. Indeed, statistical methods using time series have been developed so as to estimate parameters compatible with our model ([3, 4]). Because of the hypotheses we have made concerning birth, death and competition, our model represents a logistic population growth model with extinction. In such a setting, [4] have shown that death and birth rates can be estimated separately and so be used as parameters for our model and compare it to empirical data, either from natural or experimental populations. If our model does indeed agree with empirical observations, a natural next step would be to extend this model so as to consider multiple loci, either neutral or under selection, with possible mutation, so as to provide predictions in a more general genetic setting all the while incorporating intrinsic demographic behaviors which we may be of a great importance in shaping species diversity and evolvability.

## Acknowledgements

*This work was partially funded by the Chair “Modélisation Mathématique et Biodiversité” of VEOLIA-Ecole Polytechnique-MNHN-F.X., and was also supported by the Mission for Interdisciplinarity at CNRS and by a public grant as part of the Investissement d’avenir project, reference ANR-11-LABX-0056-LMH, LabEx LMH. Diala Abu Awad was funded by the Agence National de la Recherche (ANR SEAD - ANR-13-ADAP-0011)*.

## A Definition of *N*_*e*_ and strength of selection

Let us consider our diffusion model introduced in Equation (2)

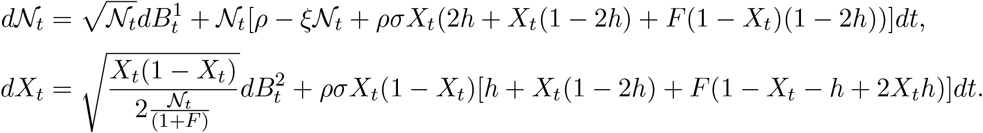

Now let us define as in [10] the time change (τ_*t*_,*t* ≥ 0) such that for all *t* > 0,

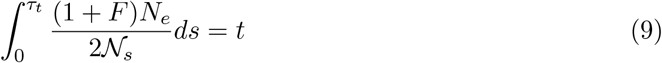

for a given real number *N*_*e*_, and let us define the time changed diffusion process 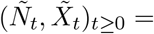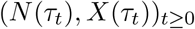.

Then 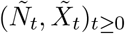 satisfies the diffusion equation:

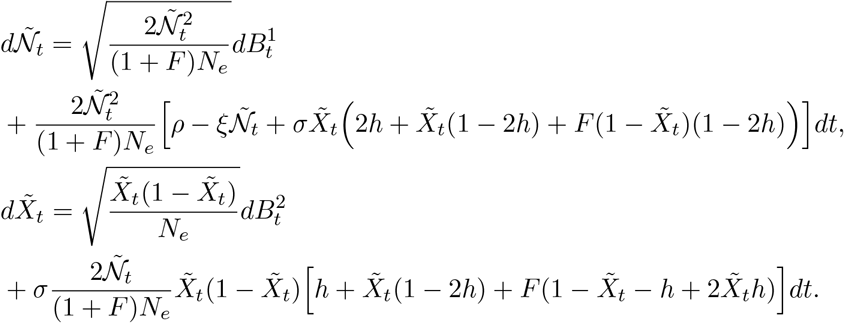

From this equation and Equation (9) we can, in a neutral case, provide a definition of the effective population mass in our model, defined as the effective population mass of a Wright-Fisher diffusion whose mean absorption time is the same than for our diffusion model with stochastically varying mass. Indeed from Equation (9)

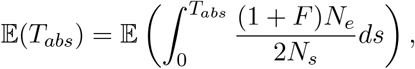

which gives,

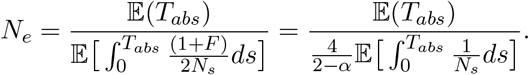

Note that using the more widely used harmonic mean of population mass so as to describe *N*_*e*_ results in over-estimations fo *N*_*e*_ (Figure A.1).

Note also that in the non-neutral case this change of time to obtain a Wright-Fisher diffusion with selection is not possible. Indeed, in this case the time-changed diffusion giving the proportion 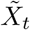 of allele *a* follows a haploid Wright-Fisher diffusion with effective population mass equal to *N*_*e*_ but with selection coefficient equal to 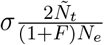 at time *t*. Population demography can therefore be seen and defined as a changing environment, though this environment is in this case itself influenced by the feedback of genetics. In this case a natural approximation is to take *s* = *σ*, as shown in Section 3.4.

**A.1:**
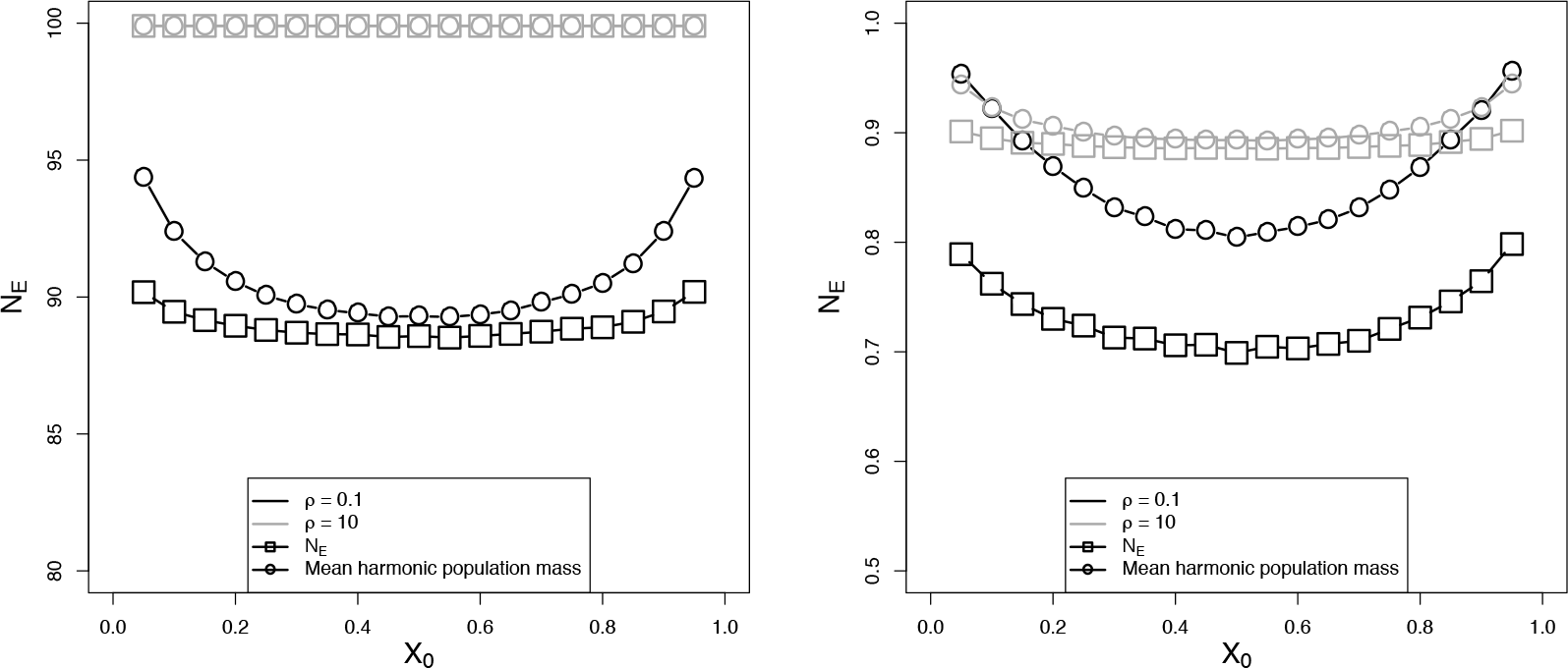
Comparing the Effective population mass proposed in equation 6 to the Mean harmonic population mass obtained from simulations run as a function of the initial frequency *X*_0_ of a neutral allele *a* (*σ* = 0) for two values of *ρ* (0.1 in black and 10 in gray) and two values of 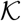, on the left 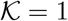 and on the right 100.

**A.2:**
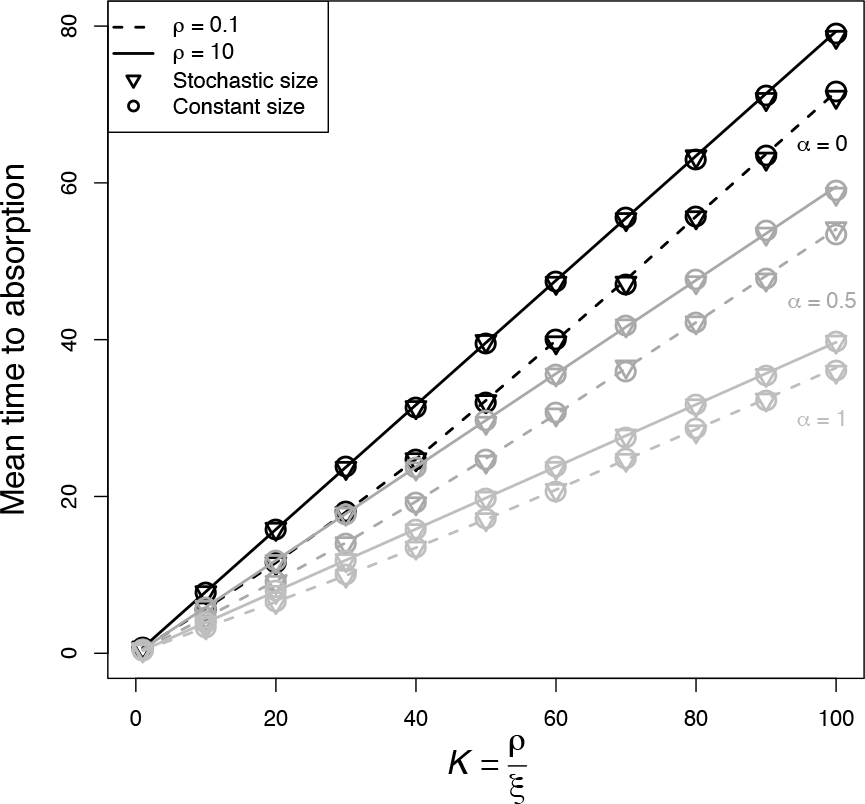
Mean times to absorption of a neutral allele (*σ* = 0) with random mating (*α* = 0), partial selfing (*α* = 0.5) and strict selfing (*α* = 1) and different values of the growth rate *ρ*, as a function of the ratio 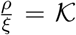 (Equation 4) for three cases: 1) Simulations of the stochastic diffusion process (2), 2) Simulations of the Wright-Fisher diffusion using *N*_*e*_ defined in Equation (6) and 3) Theoretical approximations provided by [2] using *N*_*e*_, represented by the lines (dashed for *ρ* = 0.1 nd full for *ρ* = 10.

